# Subjective confidence reveals the hierarchical nature of learning under uncertainty

**DOI:** 10.1101/256016

**Authors:** Micha Heilbron, Florent Meyniel

## Abstract

Hierarchical processing is pervasive in the brain, but its computational significance for learning in real-world conditions, with uncertainty and changes, is disputed. We show that previously proposed qualitative signatures which relied on reports of learned quantities or choices in simple experiments are insufficient to categorically distinguish hierarchical from non-hierarchical models of learning under uncertainty. Instead, we present a novel test which leverages a more complex task, whose hierarchical structure allows generalization between different statistics tracked in parallel. We use reports of confidence to quantitatively and qualitatively arbitrate between the two accounts of learning. Our results indicate that human subjects can track multiple, interdependent levels of uncertainty, and provide clear evidence for hierarchical processing, thereby challenging some influential neurocomputational accounts of learning.

## INTRODUCTION

In real-world environments, learning is made difficult by at least two types of uncertainty (Yu & Dayan, 2005). First, there is inherent uncertainty in many real-world processes. For instance, the arrival of your daily commute may not be perfectly predictable but subject to occasional delays. Faced with such random fluctuations, learners should integrate as many observations as possible in order to obtain a stable, accurate estimate of the statistics of interest (e.g. the probability of delay) (Behrens, Woolrich, Walton, & Rushworth, 2007; Yu & Cohen, 2008). Second, there is the higher-order uncertainty related to sudden changes in those very statistics (*change points*). For instance, engineering works may increase the probability of delay for an extended period. When faced with a change point, learners should discount older observations and rely on recent ones instead, in order to flexibly update their estimate (Behrens et al., 2007; Mathys, Daunizeau, Friston, & Stephan, 2011; Nassar, Wilson, Heasly, & Gold, 2010).

Confronted with both forms of uncertainty, the optimal learning strategy is to track not only the statistics of interest but also the higher-order probability of change points. This enables learners to render their estimate stable when the environment is stable (i.e. between change points) and flexible when the environment changes (Behrens et al., 2007; Iigaya, 2016; Mathys et al., 2011; McGuire, Nassar, Gold, & Kable, 2014; Meyniel, Schlunegger, & Dehaene, 2015; Payzan-LeNestour & Bossaerts, 2011). Importantly, this approach assumes that learners use a *hierarchical generative model* of their environment. Such a model comprises multiple levels, of which lower levels depend on higher ones: current observations (level 1) are generated according to statistics of observations (level 2) which themselves may undergo change points (level 3). The hierarchical approach is widely used to study learning in both health (Behrens et al., 2007; Iglesias et al., 2013) and disease (Lawson, Mathys, & Rees, 2017; Powers, Mathys, & Corlett, 2017). However, efficient learning in dynamic environments is possible *without* tracking the likelihood of individual change points (Ritz, Nassar, Frank, & Shenhav, 2017; Ryali & Yu, 2016; Sutton, 1992; Wyart & Koechlin, 2016; Yu & Cohen, 2008), and a large body of work indeed uses such a solution to model behavioral and brain responses (Bell, Summerfield, Morin, Malecek, & Ungerleider, 2016; Farashahi et al., 2017; Rescorla & Wagner, 1972). Computationally, this approach is very different as it assumes that learners do not take higher-level factors (e.g. change points) into account, and hence use a non-hierarchical or *flat* model of the world.

The possibility that the brain uses internal hierarchical models of the world is an active area of research in cognitive science (Tenenbaum, Kemp, Griffiths, & Goodman, 2011), and has important consequences for neurobiology, since hierarchical models (Friston, 2008; Lee & Mumford, 2003) and non-hierarchical ones (Farashahi et al., 2017; Yu & Cohen, 2008) require different neural architectures. In learning theory however, internal hierarchical models pose somewhat of a conundrum, being simultaneously assumed critical by some frameworks for learning under uncertainty (Behrens et al., 2007; Mathys et al., 2011; Meyniel, Schlunegger, et al., 2015) and unnecessary by others (Bell et al., 2016; Farashahi et al., 2017; Wyart & Koechlin, 2016; Yu & Cohen, 2008). One possible explanation for this conundrum is that flat approximations to hierarchical solutions can be so efficient that both accounts become difficult to distinguish. Indeed, previous studies using quantitative model comparison reported conflicting results: some authors found that learning was best explained by hierarchical models (Iglesias et al., 2013; Lawson et al., 2017; Meyniel & Dehaene, 2017; Vinckier et al., 2016) while others found that flat models best explained their results (Bell et al., 2016; Farashahi et al., 2017; Summerfield, Behrens, & Koechlin, 2011). Here, we address this issue by developing a new way to test whether learners use a hierarchical model of the world. Specifically, we sought to find a method that relied not just on comparing model fits but that can detect *qualitative signatures* or *hallmarks* that are uniquely characteristic of an internal hierarchical model.

## RESULTS

### Modulations of apparent learning rate are not a hallmark of hierarchical processing

Over and above the use of formal model comparison, earlier work on learning under uncertainty also influentially relied on demonstrations of qualitative signatures or hallmarks. In this context, a key feature of hierarchical models is the modulation of the learning rate by the changeability (or *volatility*) of the environment. One particularly influential demonstration of this principle in humans showed that the *apparent learning rate* — the ratio between the update size and the prediction error at any given observation — was modulated by change points (Behrens et al., 2007; Nassar et al., 2010). This was argued a hallmark of hierarchical processing, since increasing the learning rate after change points would only be expected from a hierarchical learner (tracking both the statistics of observations and changes in those statistics) and not from a flat learner (tracking only the statistics). However, we show a counter-example: a flat learning model whose parameters are kept fixed and nevertheless shows systematic modulations of the apparent learning rate without actually tracking the higher-level likelihood of change points (Fig 1. and Methods). Although the modulations are smaller in the flat model than in the hierarchical one, they are qualitatively identical, demonstrating that such modulations are not uniquely characteristic of hierarchical models. This suggests that the presence of apparent learning rate modulations is not sufficiently specific, and that a new test must tap into a different property in order to reveal a true hallmark of hierarchical learning.

**Figure 1.**
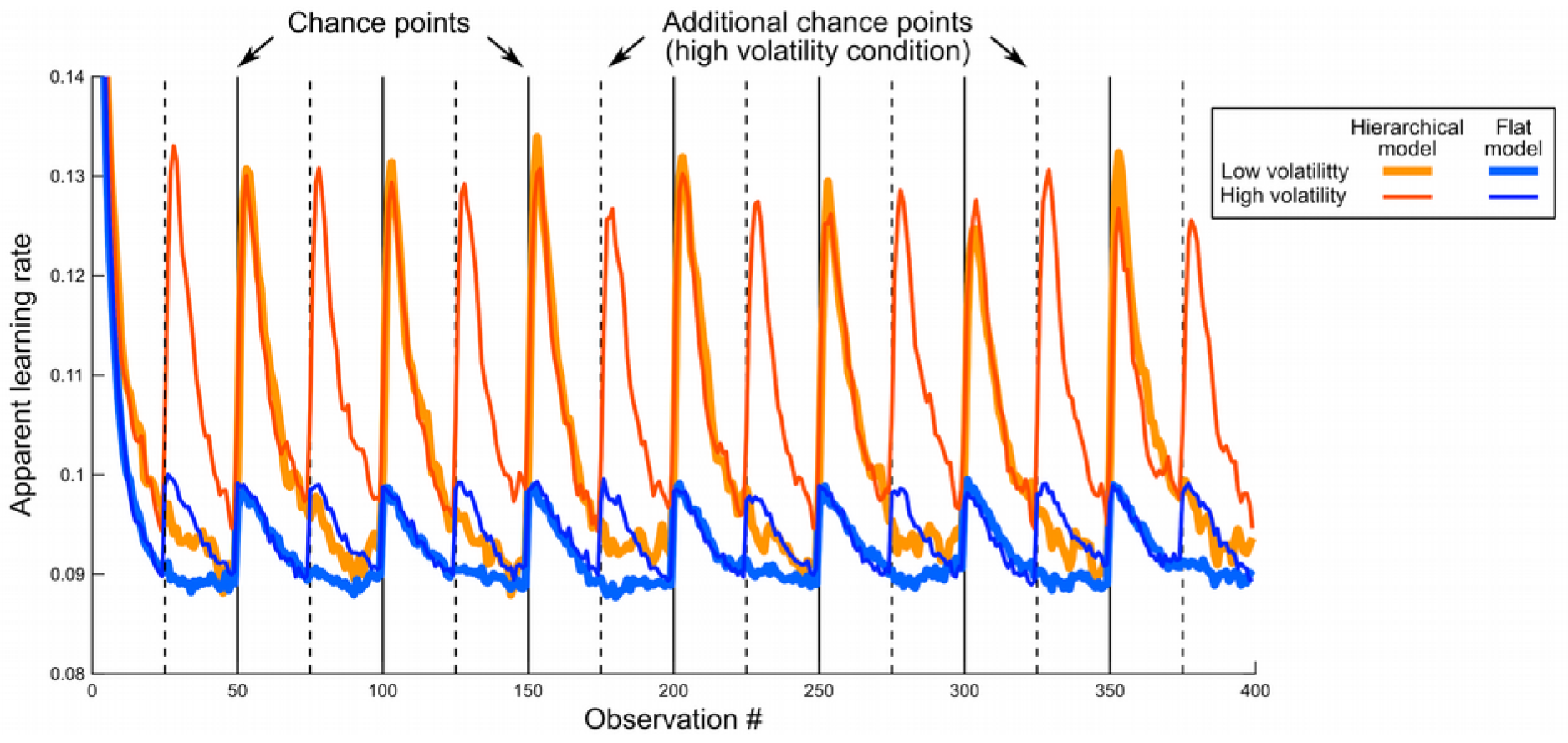
Apparent learning rate modulations in previous designs are not a hallmark of hierarchical processing. This simulation is inspired by a previous study (Behrens et al., 2007), in which subjects carried out a one-arm bandit task. The reward probability was not fixed but changed abruptly; the authors used different volatility levels (i.e. different numbers of change points). Subjects had to learn this reward probability through experience in order to optimize their payoff. Similarly, we generated sequences with low volatility (7 change points, see vertical plain black lines), and high volatility (see additional change points, vertical dashed dashed lines). The sequences were binary (absence or presence of reward) and the reward probability was resampled randomly after each change point. We consider two learning models: a hierarchical model, which estimates the reward rate, taking into account the possibility of change points; and a flat model that computes the reward rate near-optimally based on a fixed leaky count of observations, and a prior count of 1 for either outcome (see Methods). Contrary to Behrens et al, our hierarchical model does not estimate volatility, and therefore it cannot detect that the task comprises two volatilities; had we allowed for it, the observed modulations would have been even larger. Each model has a single free parameter which we fit to return the best estimate of the actual generative reward probabilities in both the low and high volatility conditions together. Keeping those best fitting parameters equal across both conditions, we measured the dynamic of the apparent learning rates of the models, defined as the ratio between the current update of the reward estimate (θ_t+1_-θ_t_) and the prediction error leading to this update (y_t+1_-θ_t_). The hierarchical model shows a transient increase in its apparent learning rate whenever a change point occurs, reflecting that it gives more weights to the observations that follow a change point. Such a dynamic adjustment of the apparent learning rate was reported in humans (Nassar et al., 2010). The flat model showed a qualitatively similar effect, despite the leakiness of its count being fixed. Note that since there are more change points in the higher volatility condition (dashed lines), the average learning rates of both models also increase overall with volatility, as previously reported in humans (Behrens et al., 2007). The lines show mean values across 1000 simulations; s.e.m. was about the line thickness and therefore omitted.

### Simulations suggest confidence offers a sensitive metric to discriminate models

When developing such a new test, the first question is what quantity or metric this test should target. In earlier studies, subjects tracked changing statistics such as the probability of a reward or a stimulus (Behrens et al., 2007; Bell et al., 2016; Gallistel, Krishan, Liu, Miller, & Latham, 2014; Iglesias et al., 2013; Jang et al., 2015; Vinckier et al., 2016), or the mean of some physical quantity like the location of a reward on an axis (McGuire et al., 2014) or its magnitude (Nassar et al., 2010). In those tasks, learning was probed either using choices that were supposedly guided by the learned statistics (Behrens et al., 2007; Bell et al., 2016; Iglesias et al., 2013; Summerfield et al., 2011; Vinckier et al., 2016) or using explicit reports of the learned statistics (Gallistel et al., 2014; McGuire et al., 2014; Meyniel, Schlunegger, et al., 2015; Nassar et al., 2010). Both choices and explicit reports are *first-order metrics*, as they only reflect the estimated statistics themselves. However, since a first-order metric only describes the level of observations, it may be seldom unique to a single model, especially if models aim at providing a good description of observations. By contrast, *second-order metrics*, such as the learner’s confidence about her estimates, describe the learner’s inference and may be more diagnostic about the underlying computations (Meyniel, Sigman, & Mainen, 2015). For illustration, we simulated a hierarchical and flat model in a probability learning problem (similar to the task used here, detailed below). Over a large range of possible task parameters, the probability estimates of the optimal hierarchical model and a near-optimal flat model were indeed highly correlated (Pearson ρ>0.9) whereas their confidence levels were much less correlated, potentially offering a more sensitive metric (see Fig. S1 and Methods).

Altogether, our simulations demonstrate firstly that modulations of the apparent learning rate are not unique to hierarchical models and are thus not a hallmark of hierarchical learning (Fig. 1); and secondly, that a new test aiming to discriminate hierarchical from flat models can use learners’ confidence about their estimates for doing so, since this metric is in theory more sensitive (Fig. S1).

### A task allowing for a more direct test for an internal hierarchical model

Based on the simulation results, we designed a new test that uses confidence to reveal clearly dissociable signatures of hierarchical and flat models of learning. Our new test builds on a task structure that has been used before (Meyniel, Schlunegger, and Dehaene 2015; Meyniel & Dehaene 2017). The motivation for using this task is that participants must track *two* changing statistics governed by the same higher-order change points. This stands in contrast to most earlier tasks discussed above, which required subjects to monitor only one changing statistic and whose hierarchical structure was therefore less prominent. Such tasks may have been too simple to categorically distinguish hierarchical and flat accounts. Here, by using multiple statistics we can probe forms of transfer *between* estimated statistics that are truly unique to internal hierarchical models, as we will detail below.

In the task (see Fig. 2A), participants observed long sequences of two stimuli (A and B), the occurrence of which was governed by two transition probabilities which subjects had to learn: p(A_t_|A_t−1_) and p(B_t_|B_t-1_). The value of each probability was independent, but at unpredictable moments (*change points*) both simultaneously changed. Subjects were fully informed about this generative process. They passively observed the stimuli and were asked to report both their estimate of the transition probability leading to the next stimulus, and their confidence in this estimate.

**Figure 2.**
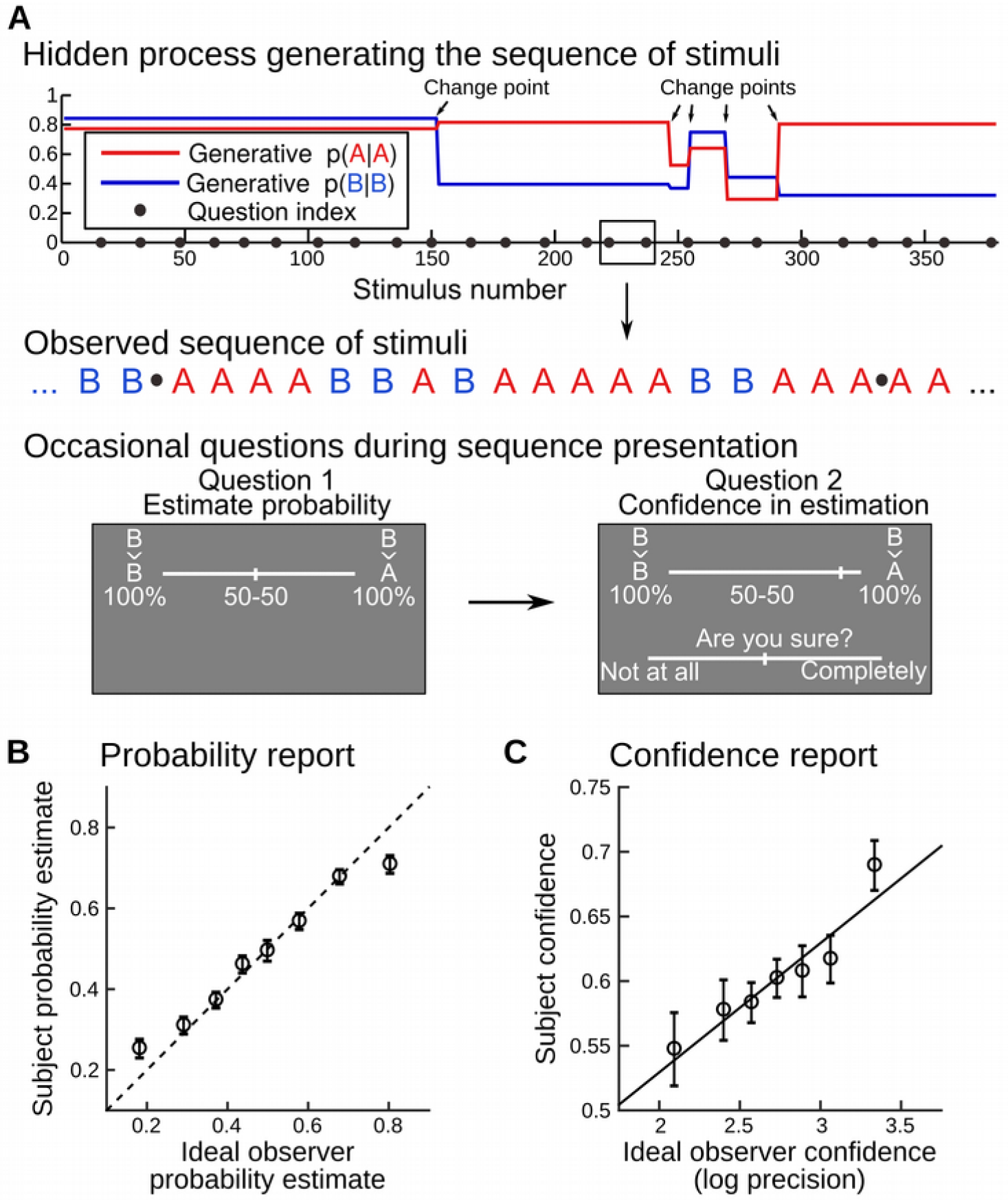
Behavioral task: learning of dynamic transition probabilities with confidence reports. (A) Probability learning task. Human subjects (N=23) were presented with random sequences of two stimuli A and B. The stimuli were, in distinct blocks, either auditory or visual and they were perceived without ambiguity. At each trial, one of either stimulus was sampled according to a probability that depended on the identity of the previous stimulus: p(A_t_|A_t-1_) and P(B_t_|B_t-1_). These transition probabilities underwent occasional, abrupt changes (change points). A change point could occur at any trial with a probability that was fixed throughout the experiment. Subjects were instructed about this generative process and had to continuously estimate the (changing) transition probabilities given the observations received. Occasionally (see black dots in **A**), we probed their inferences by asking them, first, to report the probability of the next stimulus (i.e. report their estimate of the relevant transition probability) and second, to rate their confidence in this probability estimate. (**B, C**) Subjects’ responses were compared to an ideal observer model that solved the same task usingoptimal Bayesian inference. Numeric values of confidence differ between subjects and models since they are on different scales (from 0 to 1 in the former, in log-precision unit in the latter). For illustration, the ideal observer values were binned, the dashed line (**B**) denotes the identity, the plain line (**C**) is a linear fit, and data points correspond to subjects’ mean ± s.e.m.

### Probability and confidence estimates closely follow a hierarchical ideal observer

Before testing for an internal hierarchical model, we first wanted to verify whether subjects had performed the task well, in the sense that their responses were consistent with those of an ideal observer. As a benchmark, we used the optimal ideal observer; this model was not fitted onto subjects’ data, but set so as to optimally solve the task by ‘inverting’ its hierarchical generative structure using Bayes’ rule (see methods). As displayed in Fig. 2B-C, linear regressions between participants’ responses and optimal values showed a tight agreement for probability estimates (β=0.66±0.06 s.e.m., t_22_= 11.13, p=1.7 10^−10^), and for confidence reports (β=0.10±0.03 s.e.m., t_22_= 3.06, p=0.0058); for further checks of robustness, see **Supplementary Results 2**. Despite being somewhat noisier, confidence reports also showed many properties of optimal inference (see **Supplementary Results 3**).

Since we propose that subjects’ confidence reports are more diagnostic than their first-order estimates, the next thing we verified was that confidence reports indeed conveyed information that was not already conveyed implicitly by the first-order estimates. We tested this in our data by regressing out the (theoretically expected) covariance between subjects’ confidence reports and several metrics derived from first-order estimates (see **Supplementary Results 3**); the residuals of this regression still co-varied with optimal confidence (β=0.028±0.012, t_22_=2.3, p=0.029). This result was replicated by repeating the analysis on another dataset (Meyniel, Schlunegger, et al., 2015): β=0.023±0.010, t_17_=2.2, p=0.0436; and also in the control experiment detailed below: β=0.015±0.006, t_20_=2.3, p=0.034. These results indicate that subjective confidence and probability reports are not entirely redundant, and thus that confidence is worth investigating.

Having verified that confidence and probability reports closely followed estimates of an optimal hierarchical model, and that both metrics were not redundant, we then tested whether subjects’ reports, overall, could not be better explained by a different, computationally less sophisticated model. We consider two models: the optimal hierarchical model (same as above) and a near-optimal flat model, akin to the delta-rule algorithm with a fixed learning rate (see Methods and **Supplementary Results 1**), that approximates the full Bayesian model extremely well. The models have the same number of free parameters, so model comparison boils down to comparing the goodness-of-fit. We first took the parameters that provide the best estimate of the true generative probabilities. The goodness-of-fit, assessed as mean square error (MSE) between subjects’ and models’ estimates, was better for the hierarchical model than for the flat model (paired difference of MSE, hierarchical minus flat: −0.0051±0.0014 s.e.m., t_22_=−3.7, p=0.0013). Note that subjects’ estimates of volatility, a key task parameter here, usually deviate from the optimum and show a large variability (Nassar et al., 2010; Zhang & Yu, 2013), which could bias our conclusion. We therefore fitted the model parameters per subject, and we found that the difference in fit was even more significant (−0.0077±0.0019 s.e.m., t_22_=− 3.97, p=6.5 10^−4^). This result replicates a previous finding (Meyniel & Dehaene, 2017). We then repeated the comparison for confidence levels. When model parameters were set to best estimate the true transition probabilities, the hierarchical model showed a trend toward a significantly lower MSE compared to the flat model (paired difference of MSE, hierarchical minus flat: −0.0017±0.0010 s.e.m., t_22_=−1.8, p=0.084). When model parameters were fitted onto each subjects’ confidence reports, this difference was significant (−0.0027±0.0012 s.e.m., t_22_=−2.36, p=0.028).

Altogether, these results show that participants successfully performed the task and that the hierarchical model was quantitatively superior to the flat model in explaining subjects’ probability estimates and confidence ratings. This leaves us with the last and perhaps most important question: did subjects also show a *qualitative signature* that could only be explained by a hierarchical model?

### Subjective confidence reveals a hallmark of an internal hierarchical model

Identifying the qualitative signature proposed here was possible because our task involves two transition probabilities, P(A|A) and P(B|B), whose changes were *coupled*, occurring simultaneously. In this context, a flat learner only estimates the value of each transition probability, while a hierarchical model *also* estimates the probability of a *global* change point. Faced with a global change point, the hierarchical learner then reacts optimally and makes its prior knowledge more malleable by becoming uncertain about *both* P(A|A) and P(B|B). Importantly, using this mechanism, an internal hierarchical model should allow for generalization: if a change point is suspected after observing just one type of transition (e.g. AAAAAAA, when P(A|A) was estimated to be low) a hierarchical learner would *also* become uncertain about the other quantity, P(B|B), despite having acquired no direct evidence on this transition (Fig. 3A). Critically, this form of indirect inference is unique to hierarchical models and thus offers a powerful test of hierarchical theories of learning.

**Figure 3.**
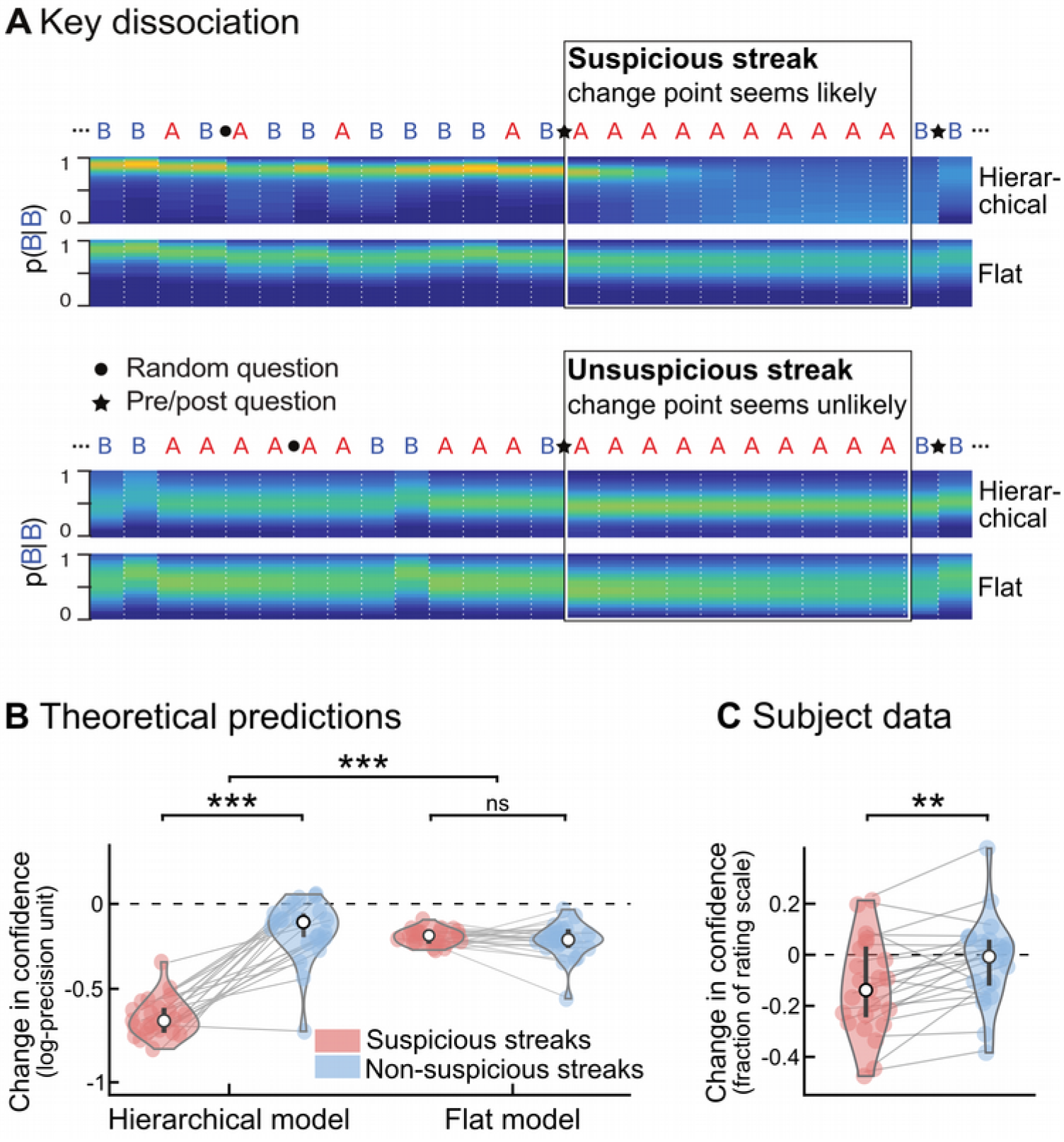
A qualitative signature of hierarchical learning in confidence reports. **(A)** Divergent predictions of hierarchical versus flat learning models. Two fragments of sequences are shown in which one stimulus (‘A’) is consecutively repeated 10 times. In the upper fragment, this streak of repetitions is highly unlikely (or ‘suspicious’) given the context, and may indicate that the underlying statistics changed. By contrast, in the lower fragment, the same streak is not unlikely, and does not suggest a change point. The heat maps show the posterior probability distribution of P(B|B), i.e. the probability of a repetition of the *other* stimulus (B), estimated by the hierarchical and flat models. In a hierarchical model, unlikely streaks arouse the suspicion of a global change in statistics, causing the model to become uncertain about its estimates of both transition probabilities, despite having acquired no direct evidence on P(B|B). In a flat model, by contrast, a suspicious streak of A’s will not similarly decrease the confidence in P(B|B), because a flat model does not track global change points. To test for this effect, pre/post questions (indicated by a star) were placed immediately before and after selected streaks, to obtain subjective estimates of the transition probability corresponding to the stimulus *not observed* during the streak. Streaks were categorized as suspicious if they aroused the suspicion of a change point from the hierarchical ideal observer’s viewpoint. Note that the flat model also shows a decrease in confidence, because it progressively forgets its estimates about P(B|B) during a streak of As, but, there is no difference between suspicious and non-suspicious streaks. **(B)** For the sequences presented to subjects, the change in confidence (post-streak minus prestreak) was significantly modulated by streak type in the hierarchical model, but not in a flat model. **(C)** Subjects’ confidence showed an effect of streak type predicted by the ideal hierarchical model. As in Fig. 2C, confidence values in subjects and models are on different scales. The error bars correspond to the inter-subject quartiles, distributions show subjects’ data; significance levels correspond to paired t-test with p<0.005 (**) and p<10^−12^ (***).

To test for this generalization effect, we focussed on streaks of repetitions, and distinguished between streaks that seem unlikely in context and may signal a change point (*suspicious streaks*) and streaks that do not (*non-suspicious streaks*). Stimulus sequences were carefully selected to contain enough suspicious and non-suspicious streaks and to control for confounds such as streak duration (see Methods). Questions were inserted just before and after the streak, so that subjects reported their estimate of (and confidence in) the other, non-repeating transition (Fig. 3A). Exact theoretical predictions for both models are found in Fig. 3B. In the the hierarchical model, confidence decreases strongly after suspicious, but much less after non-suspicious streaks. In the flat model, however, there is no such difference. Strikingly, subjective reports exactly followed the hierarchical account and falsified the flat one, see Fig. 3C: confidence decreased strongly after suspicious (−0.12±0.04 s.e.m, t_22_=−3.2, p=0.004) but not after non-suspicious streaks (−0.02±0.03 s.e.m., t_22_=−0.7, p=0.51), and this interaction was significant (paired difference, 0.10±0.03 s.e.m., t_22_=3.7, p=0.001).

### Various controls demonstrate the specificity of the effect on confidence

We now rule out a series of potential confounding explanations. First, one concern is that the analysis above involves models that are optimized to estimate the true transition probabilities of the task: perhaps the predictions look different if we fit the models onto behaviour. However, our conclusions remain unaffected if we simulate models fitted onto each subject, (Fig. S2 C, E). Another concern is that our analyses assume subjects were tracking transition probabilities, while they may in fact have been tracking another (heuristic) quantity, perhaps using a flat model. Detailed analysis revealed that subjects did in fact track transition probabilities (see **Supplementary Results 3**) and that no heuristic flat model could explain the selective decrease of confidence (Fig S2 B, D, F).

One may also wonder whether the effect reported in Fig. 3 about confidence is also found in another variable. Note that this would not invalidate our conclusion about the hierarchical nature of learning. Fig. S3 shows that probability estimates (the ones about which confidence is reported and shown in Fig. 3) are not affected by streak types neither in subjects (paired difference between streak types, −0.01±− 0.02 s.e.m., t_22_=−0.5, p=0.59) nor in the hierarchical model (−0.01±0.01 s.e.m., t_22_=−1.4, p=0.17). A more subtle effect is that, when a change point is suspected, generalization should lead to reset the estimate of the unobserved transition probability, which should thus get closer to the prior value 0.5. However, this effect is less straightforward, because the estimated transition probability may already be close to 0.5 before the streak, such that an effect of streak type on the distance to the prior may be difficult to detect. Indeed, in the hierarchical model, streak type had only a weak effect (paired difference, 0.02±0.01 sem, t_22_=2.9, p=0.008). For comparison, the same effect on confidence (Fig. 3) had t_22_=11.7, p=6.9 10^−11^. The expectedly weaker effect of streak type on the distance to the prior was not detected in participants (−0.0036 +/− 0.01 sem, t_22_=−0.3, p=0.76). We also tested reaction times since they often co-vary with confidence. Here, when the optimal confidence was lower, subjects took longer to respond to the prediction question (slope of reaction times vs. optimal confidence: −0.57±0.19 s.e.m., t_22_=−3.07, p=0.005), but not to the confidence question (slope: 0. 04±0.08 sem, t_22_=0.48, p=0.64). However, there was no effect of streak type on reaction times both for the probability estimate and reports (paired difference between streak types, both p>0.27).

A final alternative explanation for the effect shown in Fig. 3 is that suspicious streaks were more surprising and that subjects may become *generally uncertain* after surprising events. In this case, the effect would not reflect hierarchical inference but simply general surprise. We therefore performed a control experiment in which both probabilities changed independently: here, suspicious streaks were equally surprising but no longer signalled a *global* change point (Fig. 4A). Indeed, generalization of a decrease in confidence was no longer observed for the hierarchical model or in subjects (paired difference between suspicious and non-suspicious streaks: 0.03±0.02 s.e.m., t_20_=1.5, p=0.15), see Fig. 4B. This absence of effect in the control task is significantly different from the effect found in the main task (difference of paired differences, two-sample t-test, t_42_=−2.03, p=0.048). This difference is not due poor performance in the control experiment (see Fig 4C): linear regression between the optimal hierarchical model for uncoupled change (the ideal observer in this task) and subjects showed a tight agreement for both predictions (β=0.61±0.06 s.e.m., t_20_= 10.27, p=2 10^−9^) and confidence (β=0.08±0.01 s.e.m., t_20_= 6.83, p=1.2 10^−6^), as in the main task (see Fig. 1 B, C). The difference between the two tasks shows an effect of higher-level factors (coupled vs. uncoupled change points) and thus constitutes further evidence for a hierarchical model. Altogether, learners generalize from inferring the change of one probability to decreasing their confidence in the estimate of another probability, but only when they know the changes are coupled and it is thus adaptive to do so. This supports that the result in Fig. 3 and **4** is a hallmark of an internal hierarchical model, and does not reflect a simpler, heuristic inference.

**Figure 4.**
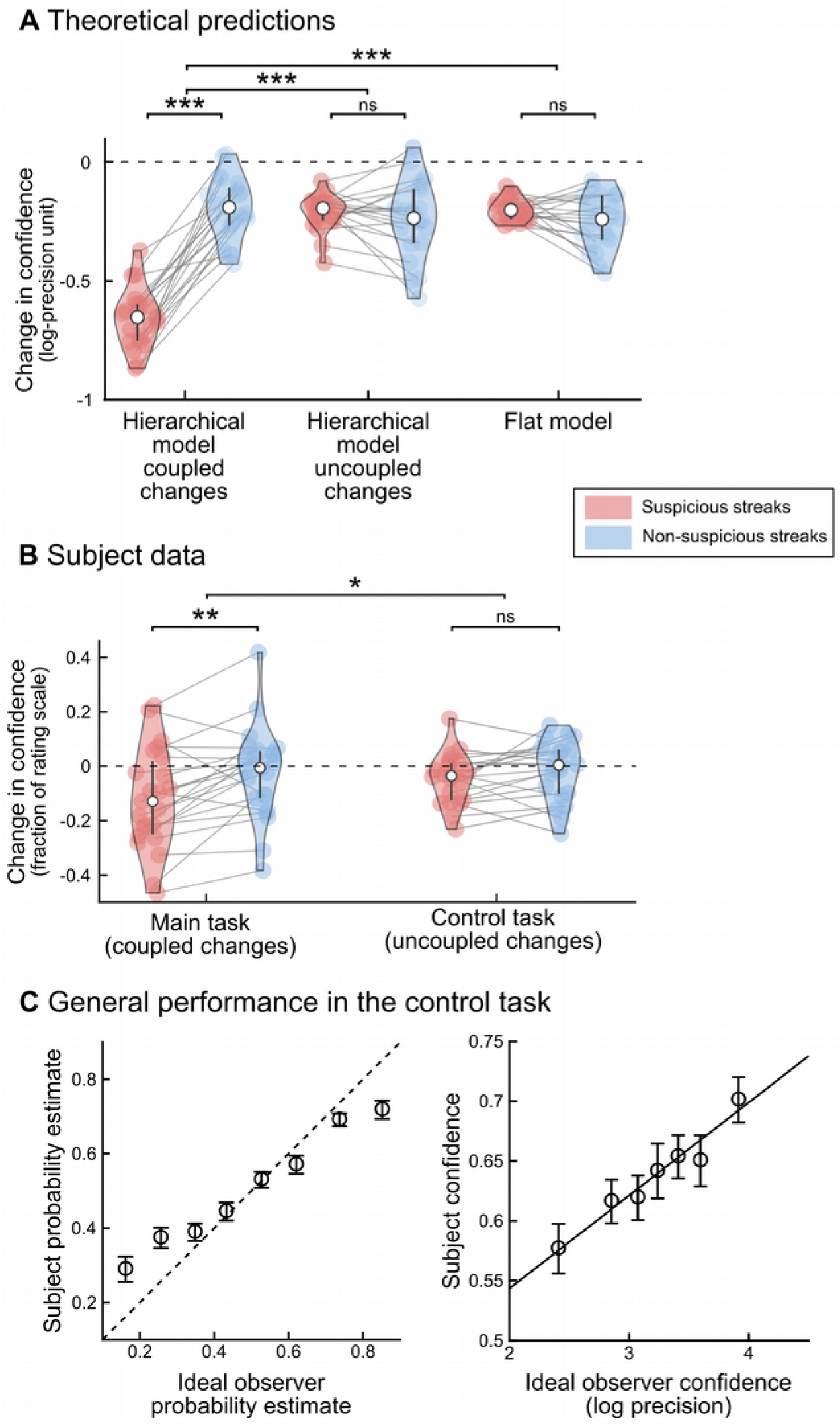
Control experiment: subjects take into account the higher-order structure of the dynamics. Results of the control experiment, in which change points were uncoupled between the two transition probabilities, thereby abolishing the possibility to infer a change in one transition probability by only observing the other transition type. (**A**) Theoretical predictions for changes in confidence around the target streaks. The optimal hierarchical model for the main task assumes that change points are coupled (“hierarchical model, coupled changes”), which is no longer optimal in the case of uncoupled change points. This model was nevertheless used to identify suspicious and non-suspicious streaks and indeed it showed an effect of streak type on the change in confidence here inthe control task as in the main task (Fig. 3C). The optimal hierarchical Bayesian model for this control experiment is similar to this first model, the only difference is that it assumes that change points are uncoupled (“hierarchical model, uncoupled changes”). As expected, this model correctly showed no effect of streak type on the change in confidence. The flat model, by definition, ignores change points and therefore whether they are coupled or uncoupled, as a result it shows no effect of streak type (as in the main experiment). (**B**) Subjects showed no difference between streak types, like the hierarchical model for uncoupled changes. The results of the main task are reproduced from Fig. 3C to facilitate visual comparison. (**C**) Subjects overall perform well in the control task, showing a tight agreement with the optimal hierarchical model for uncoupled change (the optimal ideal observer in this task) for both predictions (left) and confidence (right). In panels **A** and **B**, the error bars correspond to the inter-subject quartiles, distributions show subjects’ data. In panel **C**, data points are mean ± s.e.m across subjects. In all panels; significance levels correspond to p<0.05 (*), p<0.01 (**), p<0.001(***) in a t-test.

## DISCUSSION

We have shown that some previous tests are insufficient to categorically distinguish hierarchical from non-hierarchical models of learning in uncertain and changing environments, and we introduced a novel test to dissociate the two. The key features of our experiment are that subjects estimate two statistics that depend upon the same change points, and that we analyse the subjects’ confidence about their estimation. Our test taps into a unique property of hierarchical models: the ability to generalise between different probabilities that are coupled by the higher-order structure of the task. As such, both classes of theories make *qualitatively different predictions* at the level of individual trials. Based on both qualitative and quantitative model comparison, our results provide clear evidence in support of a hierarchical account of adaptive learning in humans. These results indicate that humans can track multiple levels of uncertainty that are hierarchically organized, including the uncertainty about their own inference.

### Quantitative vs. qualitative model comparison

A widely used method for contrasting competing models is quantitative model comparison. Following this approach, multiple models are fit onto the subjects’ data, and the model that achieves the best fit with respect to its complexity is deemed most likely (Gelman et al., 2013; Stephan, Penny, Daunizeau, Moran, & Friston, 2009). This approach is attractive because it is generally applicable and it provides a common metric (e.g. goodness-of-fit, Bayes-Factor, exceedance probability) to compare different models. This approach nevertheless needs to be supplemented for at least two reasons. First, quantitative model comparison only allows for relative conclusions, such as one model being *better than the other tested models*, but it does not allow for more general conclusions such as the *falsification* of one model (Palminteri, Wyart, and Koechlin 2017). Moreover, it is not always clear what underlying factors are contributing to differences in the models’ goodness-of-fit, and whether these factors are indeed most relevant to the question at hand. A complimentary approach, is to analyse specific, critical trials for the presence of *qualitative signatures* or *hallmarks* that uniquely identify or exclude one type of model. Such signatures are appealing because they are easily interpretable and they directly reflect a critical theoretical distinction. Previous prominent examples of the use of qualitative tests include the influential two-step task used to distinguish model-based from model-free reinforcement learning (Daw, Gershman, Seymour, Dayan, & Dolan, 2011). Here, both approaches are indeed complementary: quantitative model comparison provided evidence in favor of a hierarchical account of learning, and the qualitative approach tested for unique hallmarks and thereby falsified a flat account.

### Which behavioral metrics best reveal the learner’s computations?

A feature that distinguishes our task from previous studies is the use of explicit confidence ratings to test a key dissociation between flat and hierarchical models (Fig. 3). Our rationale was that since the flat models considered here are known to provide very accurate first-order approximations to hierarchical (optimal) models (Meyniel, Maheu, & Dehaene, 2016; Sutton, 1992; Yu & Cohen, 2008), we should opt for another variable that is less correlated between models. Choices and first-order reports are often used in behavioral science, but other metrics like subject’s confidence and reaction times also proved useful to study cognition, see (Shadlen & Kiani, 2013). Here, our simulations showed that confidence is less correlated between models, we therefore took confidence as a window on the learner’s computations.

In principle it should be possible to use other metrics to probe the same effect. Reaction times are one obvious candidate, inasmuch they are often an implicit measure of the subject’s uncertainty (Dotan, Meyniel, & Dehaene, 2017; Kepecs & Mainen, 2012; Kepecs, Uchida, Zariwala, & Mainen, 2008; Kiani, Corthell, & Shadlen, 2014). In our study, we found a correlation between reaction times and confidence, but no effect of streak type that serves as our hallmark signature. In our study, in contrast to accumulation (Kiani et al., 2014) or waiting-time (Kepecs et al., 2008) paradigms, there is no principled reason for reaction times to co-vary with confidence, which may explain why reaction times do not show the hallmark signature of a hierarchical inference.

Another candidate is the *apparent learning rate*. Previous studies reported modulations of the apparent learning rate by change points (Behrens et al., 2007; Nassar et al., 2010). The optimal, hierarchical model indeed shows such modulations because its updates are confidence-weighted (Mathys et al., 2011; Meyniel & Dehaene, 2017): for a given prediction error, its updates are larger when confidence about prior estimates is lower, which is typically the case when a change point is suspected. However, we found that in simple experiments that require to monitor only the frequency of a stimulus or a reward, a flat model could exhibit similar modulations, which are therefore not diagnostic of a hierarchical inference. In more complex experiments like the one here, the apparent learning rate could nevertheless show our hallmark signature of a hierarchical inference. Our theoretical analysis supports this hypothesis (see **Supplementary Results 4**) but we cannot assess it in our data, since this analysis requires a trial-by-trial measure of the apparent learning rate, and thus trial-by-trial (not occasional) reports of first-order estimates. A trial-by-trial measure of the apparent learning rate is neither accessible if subjects make choices at each trial. In such studies (Behrens et al., 2007; Glaze, Kable, & Gold, 2015), the authors could only use choices to compute an apparent learning rate in a sliding window of trials but this analysis lacks the trial-by-trial resolution. In our task, one could investigate the apparent learning rate of subjects, but that would require subjects to report their probability estimates after each trial, and hence to constantly interrupt the stimulus stream. This would probably interfere with the participants’ ability to integrate consecutive observations, which is critical for tracking transition probabilities, and therefore seems difficult to implement in practice. Furthermore, if an effect of streak type were observed on the apparent learning rate, it would probably be mediated by the subject’s confidence (Mathys et al., 2011; Meyniel & Dehaene, 2017), in that case one may prefer to probe confidence directly.

We acknowledge that there are drawbacks of using confidence as the metric of interest. In theory, confidence more reliably discriminates flat and hierarchical models; but in practice we found that the agreement between participants and the ideal observer was considerably more precise for probability estimates than for confidence ratings (see Fig. 3B-C and Fig 4C). This noisy character of confidence measurements was also reported previously (Baranski & Petrusic, 1994; Maniscalco & Lau, 2012; Meyniel, Schlunegger, et al., 2015; Zylberberg, Barttfeld, & Sigman, 2012), it may hinder the use of confidence as a metric to discriminate between models, and it may explain that the difference in overall model fit between the flat and hierarchical model here was weaker for confidence than for probability estimates. This problem may be even worse when using an indirect indicator of confidence, such as reaction times or the apparent learning rate.

### Learning in a structured environment

Our qualitative test for a hierarchical inference also leverages a particular task structure: the higher-level dependence of generative statistics upon the same (or distinct, in the control experiment) change points. Our task structure is more complex than experiments that require to monitor only one generative statistics (Behrens et al., 2007; Gallistel et al., 2014; Iglesias et al., 2013; Kheifets & Gallistel, 2012; McGuire et al., 2014; Nassar et al., 2010). Our task structure may constrain the applicability and generality of our experiment, but it has a certain ecological appeal since in real-life situations, multiple regularities are often embedded in a single context. We believe that more complex task structures are suited to distinguish complex computations and approximations. Both are likely to be equivalent in simpler experiments, whereas in highly structured environments with multiple interdependent levels (Schapiro, Rogers, Cordova, Turk-Browne, & Botvinick, 2013; Tenenbaum et al., 2011), an optimal learning algorithm can hardly obliviate the hierarchical nature of the problem to solve.

An interesting and difficult problem that we leave unaddressed here is how subjects may discover the task structure (Pouget, Beck, Ma, & Latham, 2013; Tenenbaum et al., 2011; Tervo, Tenenbaum, & Gershman, 2016). In our task, the optimal hierarchical model is able to correctly identify the current task structure (coupled vs. uncoupled change points), but only with moderate certainty even after observing the entire experiment presented to one subject (log-likelihood ratios range from 2 to 5 depending on subjects). Therefore, in principle, subjects who are not endowed with optimal computing power cannot identify reliably the correct structure from observations alone. We speculate that in real-life situations, some cues or priors inform subjects about the relevant dependencies in their environment; if true, then our experiment in which subjects were instructed about the correct task structure may have some ecological validity.

Interestingly, while the importance of hierarchical inference remains controversial in the learning literature (Bell et al., 2016; Farashahi et al., 2017; Gallistel et al., 2014; Iglesias et al., 2013; Mathys et al., 2011; McGuire et al., 2014; Nassar et al., 2010; Ritz et al., 2017; Ryali & Yu, 2016; Summerfield et al., 2011; Wyart & Koechlin, 2016), it seems more clearly established in the domain of decision making and action planning (Balaguer, Spiers, Hassabis, & Summerfield, 2016; Daw et al., 2011; Huys et al., 2012; Keramati, Smittenaar, Dolan, & Dayan, 2016; Schapiro et al., 2013; Wunderlich, Dayan, & Dolan, 2012). For instance, it was suggested that the functional organization of cognitive control is nested: low level cues trigger particular actions, depending on a stimulus-response association which is itself selected depending on a particular context (Koechlin, Ody, & Kouneiher, 2003). In this view, negative outcomes may indicate that the (higher-level) context has changed and thus that a new rule now applies. This inference even seems to be confidence-weighted in humans: the suspicion of a change in context is all the stronger that subjects were confident that their action should have yielded a positive outcome under the previous context (Purcell & Kiani, 2016). Those two studies feature an important aspect of hierarchy: a succession of (higher-level) task contexts separated by change points governs the (lower-level) stimuli. Our task also leverages another feature of hierarchy: it allows generalization and transfer of knowledge. A rule learned in a particular context can be applied in other contexts, for instance see (Collins, Cavanagh, & Frank, 2014; Collins & Frank, 2016). Our results go beyond a mere transfer: they show that the brain can *update* a statistics in the absence of direct evidence thanks to higher-level dependencies.

### Possible neural implementation for adaptive learning

Our results falsify a flat account of learning in our task, therefore they also falsify the possible use of a simple leaky integration by neural networks to solve this task. Leaky integration is often deem both plausible biologically and computationally efficient (Farashahi et al., 2017; Glaze et al., 2015; Rescorla & Wagner, 1972; Yu & Cohen, 2008). A sophisticated version of the leaky integration with metaplastic synapses allows partial modulation of the apparent learning of the network, without tracking change points or volatility (Farashahi et al., 2017). Others have suggested that computational noise itself could enable a flat inference to automatically adapt to volatility (Wyart & Koechlin, 2016). Those approximate solutions dismiss the need to compute higher-level factors like volatility, they are thus appealing due to their simplicity; however, we believe that such solutions cannot explain the generalization afforded by hierarchical inference that we showed here. We nevertheless acknowledge that it is unlikely that only one algorithm subserves all forms of learning in the brain, and therefore that our result does not dismiss the possibility that the brain resorts to flat, simpler and yet efficient algorithms like the delta rule in many situations. One previously proposed bio-inspired model seems compatible with our result (Iigaya, 2016). This model comprises two modules: one for learning and the other for detecting change points, or “unexpected surprise” (Yu & Dayan, 2005). When a change point is detected, a reset signal is sent to the learning module. Converging evidence indicates that noradrenaline could play such a role (Bouret & Sara, 2005; Nieuwenhuis, Aston-Jones, & Cohen, 2005; Salgado, Treviño, & Atzori, 2016; Schomaker & Meeter, 2015). A global reset signal could promote learning for the two transition probabilities that are maintained in parallel in our task, thereby allowing the reset of both when only one arouses the suspicion of a change point. Such a hypothesis nevertheless needs to be refined in order to account for the fact that the two statistics can also be reset independently from one another, as in the control task.

We hope that the test which we propose for hierarchy here will be applied to other learning model that computes uncertainty, and even to non-human animals despite significant methodological challenges, and therefore that it will be of interest to experimentalists and theoreticians alike.

## MATERIALS AND METHODS

### Participants

Participants were recruited by public advertisement. They gave a written informed consent prior to participating and received 20 euros for volunteering in the experiment. The study was approved by the local Ethics Committee (CPP n°08–021 Ile de France VII). 26 participants (17 female, mean age 23.8, s.e.m.: 0.49) performed the main task and 21 other participants performed the control task (11 female, mean age 23.0, s.e.m.: 0.59). We excluded participants who showed poor learning performance, which we quantified as the Pearson ρ coefficient between their probability estimates and the ideal observer’s estimates. We used a threshold corresponding to 5% of the (lowest) values measured in this task (ρ<0.18, from a total of 105 participants in this study and others) This excluded 3 subjects from the main task, and none from the control task.

### Main Task

The task was run using Octave (Version 3.4.2) and PsychToolBox (Version 3.0.11). Each participant completed a total of 5 blocks: 1 training block and 4 experimental blocks (2 auditory, 2 visual). Auditory and visual blocks alternated, with the modality of the first block randomised across participants. In each block, we presented binary sequences of 380 stimuli (1520 total) denoted A and B, which were either visual symbols or sounds and were perceived without ambiguity.

Sequences were generated according to the same principles as in previous studies (Meyniel & Dehaene, 2017; Meyniel, Schlunegger, et al., 2015). A and B were randomly drawn based on two hidden transition probabilities which subjects had to learn. These probabilities were stable only for a limited time. The length of stable periods was randomly sampled from a geometric distribution with average length of 75 stimuli, truncated at 300 stimuli to avoid overly long stable periods. Critically, and contrary to other studies (Behrens et al., 2007) the volatility was thus fixed (at 1/75). Transition probabilities were sampled independently and uniformly between 0.1-0.9, with the constraint that, for at least one of the two probabilities, the change in odd ratio (p/1-p) between consecutive stable periods was at least fourfold, thus guaranteeing that the change was effective. Across sequences and subjects, the actually used generative values indeed covered the transition probability matrix 0.1-0.9 uniformly, without any correlation (Pearson ρ = −0.0009, p = 0.98). Occasionally, the sequence was interrupted and subjects had to estimate the probability that the next stimulus would be either an A or a B and report their confidence in that estimate. Questions were located quasi-randomly, semi-periodically once each 15 stimuli on average (100 in total). Of the 100 questions, 68 questions were randomly placed; the remaining 32 questions were intentionally located just before and after 16 selected streaks (8 suspicious, 8 non-suspicious) and functioned as pre/post-questions to evaluate the effect of these streaks (see Fig. 3). For details on the definition and selection of suspicious/non-suspicious streaks, see below.

To familiarize participants with the task they were carefully instructed and performed one training block of 380 stimuli (or ~12 minutes). To make sure they were fully aware of the volatile nature of the generative process, participants had to report when they detected changes in the hidden regularities. In the experimental blocks, reporting change points was omitted, but participants knew the underlying generative process was the same.

### Control Task

The control task was very similar to the main one, with only two differences. (1) When a change occurred, it impacted only one of the two transition probabilities (randomly chosen). (2) During the training block, when subjects were required to report when they detected change points, they also reported which of the two transition probabilities had changed.

### Selection of sequences

Each randomly generated sequence was evaluated computationally and carefully selected to ensure that each subject encountered enough target moments during which the models make qualitatively different predictions, and that all sequences were balanced in terms of potential confounds such as streak duration and location. To this end, 4 random sequences of 380 stimuli long (each corresponding to one block) were analyzed computationally with the hierarchical and flat learning models, yielding 4 simulated ‘blocks’. The sequences, and associated trial-by-trial transition probability estimates from both models, were concatenated to form a single experimental sequence (of 1520 stimuli). This experimental sequence was then submitted to several selection criteria. First, we assessed whether the sequence contained at least 8 suspicious and 8 non-suspicious ‘streaks’. Consecutive repetitions were defined as ‘streaks’ if they consisted of at least 7 or more stimuli, and started after the 15th stimulus of a block. Streaks were classified as ‘suspicious’ if they aroused the suspicion of a change in the hierarchical ideal observer. Computationally, this was defined via the confidence in the probability of the observed repetition decreasing at least once during the streak. Following this criterion, even streaks of repetitions are that just slightly surprising are considered ‘suspicious’. To ensure the effect would be observable, only sequences in which the suspicious streaks led to a sizeable decrease in theoretical confidence levels were selected. Due to an error in the selection procedure, one sequence was included for which the theoretically expected average decrease in confidence after non-suspicious streaks was in fact larger than that after suspicious streaks. Because the corresponding subject who observed this sequence did show sufficient learning performance and hence added valuable data to all other analyses, we decided not to exclude the participant from the study. Importantly, excluding this subject does not change any conclusion or significance level of the statistical tests reported here.

To control for factors that may potentially confound decreases in confidence, only sequences in which the average duration of suspicious and non-suspicious streaks was approximately identical, and in which there was at least one streak of each type in each block, were selected. In addition, subjects were not informed about the distinction between suspicious and non-suspicious streaks or that between random questions and pre-post questions that targeted the critical moments before and after streaks. Interviews performed after the experiment ruled out that subjects understood the goal of the experiment, as no subject had noticed that a sizable fraction (~30%) of questions purposefully targeted streaks.

### Ideal observer models

The models used in this study are implemented in a Matlab toolbox available on GitHub and described in a previous publication (Meyniel et al., 2016). The model termed “hierarchical” and “flat” here correspond respectively to the hidden Markov model (HMM) and the leaky integrator model in the toolbox. Here, we summarize the essential aspects of those models.

The hierarchical and flat models (M) are both ideal observer models: they use Bayes rule to estimate the posterior distribution of the statistic they estimate, *θ*_*t*_, based on a prior on this statistic and the likelihood provided by previous observations, *y*_*1:t*_ (here, a sequence of As and Bs):

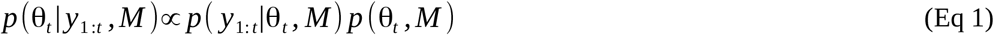

Subscripts denote the observation number within a sequence. In the main text, the models estimate the transition probabilities between successive stimuli, so that θ is a vector with two elements: θ = [p(A|A), p(B|B)]. Note that those two probabilities suffice to describe all transitions, since the others can be derived as p(B|A) = 1-p(A|A) and p(A|B)=1-p(B|B). In Fig. S2, we also consider variants in which the model estimate another statistic, the frequency of stimuli: θ = p(A). Note that p(B) is simply 1-p(A).

The estimation of θ depends on the assumption of the ideal observer model (M). The flat model considers that θ is fixed, and evaluates its value based on a leaky count of observations. The internal representation of this model therefore has only one level: θ, the statistic of observations. When the true generative statistic is in fact changing over time, the leakiness of the model enables it to constantly adapt its estimate of the statistic and therefore to cope with changes. If the leakiness is tuned to the rate of change, the estimate can approach optimality (see Fig. S1A).

By contrast, the hierarchical model entertains the assumption that θ can abruptly change at any moment. The internal representation of the model therefore has several levels beyond observations: a level characterizing the statistic of observations at a given moment (θ_t_) and a level describing the probability that of a change in θ occurs (p_c_). Conceivably, there could be higher-order levels describing changes in p_c_ itself (Behrens et al., 2007); however this sophistication is unnecessary here and we consider that p_c_ is fixed.

#### Flat model

The flat model assumes that the true value of θ is fixed, and it constantly infers its value given the evidence received. Therefore, the likelihood function can be decomposed as follows:

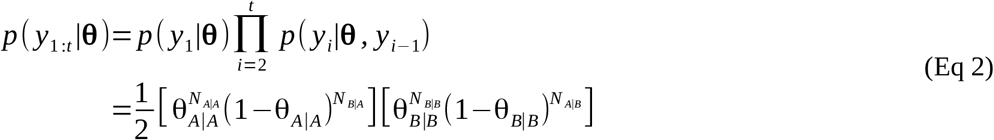

Where *N*_A|A_(t) denotes the number of AA pairs in the sequence *y*_1:t_. A convenient parametrization for the prior distribution is the beta distribution: 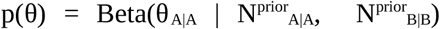. This parametrization allows for an intuitive interpretation of N^prior^_A|A_ and N^prior^_B|B_ as prior observation counts, and due to its conjugacy with the likelihood function (Eq2), inserting Eq2 into Eq1 yields that the posterior probability of θ is the product of two beta functions:

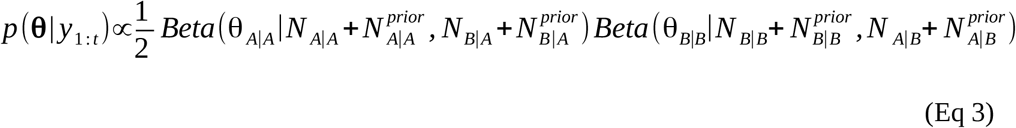

We consider here that the count of observations (the number of AA and BB pairs) is leaky, so that observations that are further in the past have a lower weight than recent ones. We modeled this leakiness as an exponential decay ω, such that the k-th past stimulus has a weight e^−k/ω^. Note a perfect integration, in which all observations are given the same weight, corresponds to the special case with ω being infinitely large. Also note that ω*ln(2) corresponds to the “half-life”, i.e. the number of observations after which the weight of a past observation is reduced by half.

In the main text (but Fig. 1), we choose [N^prior^_A|A_, N^prior^_B|B_] = [0 0], for in this case, the mean estimate of the flat model becomes strictly equivalent to the estimate of a “delta rule” as the number of observations increases (see **Supplementary Result 1**). An alternative choice for the prior is the so-called Laplace-Bayes prior [1 1], which is uninformative in that it gives the same prior probability to any value of θ (Gelman et al., 2013). This choice is important for Fig. 1, but not for the results in the main text (see Fig S2).

#### Hierarchical model

The hierarchical model evaluates the current value of the generative statistic θ under the assumption that it may change at any new observation with a fixed probability p_c_. Note that, would the location of the change points be known, the inference of θ would be simple: one would simply need to count the number of pairs (*N*_A|A_, *N*_B|B_, *N*_A|B_, *N*_B|A_) since the last change point and apply Eq. 2. However, without knowing the location of change points, one should in principle average the estimates given all possible locations of change points, which is in practice far too large a number. The computation is rendered tractable by the so-called Markov property of the generative process. If one knows θ at time *t*, then the next observation *y*_*t*+1_ is generated with θ_*t*+1_ = θ_*t*_ if no change occurred and with another value drawn from the prior distribution otherwise. Therefore, if one knows θ_*t*_, previous observations are not needed to estimate θ_*t*+1_. Casting the generative process as a Hidden Markov Model (HMM) enables to compute the joint distribution of θ and observations iteratively, starting from the prior, and updating this distribution by moving forward in the sequence of observations:

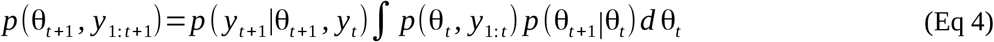

This integral can be computed numerically by discretization on a grid. The posterior probability can be obtained by normalizing this joint distribution.

#### Probability reports and confidence ratings with the models

Both the flat and hierarchical models estimate a full posterior distribution for θ, therefore both models have a posterior uncertainty (or conversely, confidence) about their estimate. In that sense, the flat model can be considered as a delta rule that is extended to provide confidence estimates about first-order estimates (see **Supplementary Results 1** for more details about the flat model and delta rule).

The probability of the next stimulus (question #1 asked to subjects) was computed from the posterior using Bayes rule:

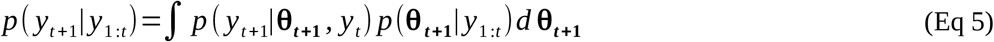

Note that the first term in the integral, the likelihood, is nothing but the relevant transition probability itself (conditioned on the actual previous observation). This integral is therefore simply the mean of the posterior distribution of the relevant transition probability. The confidence in the reported probability estimate (question #2) was computed as the log-precision of this posterior distribution (Meyniel & Dehaene, 2017; Meyniel, Schlunegger, et al., 2015; Meyniel, Sigman, et al., 2015).

### Model fit

The flat and the hierarchical models have one free parameter each, respectively ω (the leakiness) and p_c_ (the prior probability of change point).

Unless stated otherwise, the analysis reported in the main text used the parameters that best fit the true probabilities used to generate the sequences of observations presented to subjects. More precisely, for each sequence of observations, we computed the probability of each new observation given the previous ones, as estimated by the models using Eq. 5 and we compared it to the true generative probability. We adjusted the free parameters ω and p_c_ with grid-search to minimize the sum of squared errors (SSE) over all the sequences used for all subjects. The resulting values, ω=20.3 and p_c_=0.014 (indeed close to the generative value 1/75).

We also fitted the parameters to the responses of each subject (Fig. S2). For probability estimates, the above grid-search procedure was repeated after replacing generative values with the subject’s estimates of probabilities at the moment of questions. For confidence reports, we used a similar procedure; note however that subjects used a bounded qualitative slider to report confidence whereas the model confidence is numeric and unbounded, so that there is not a direct mapping between the two. Therefore, the SSE was computed with the residuals of a linear regression between subject’s confidence and the model’s confidence.

### Statistical analyses

All linear regressions between dependent variables (e.g. probability estimates, confidence ratings) and explanatory variables (optimal estimates of probabilities and confidence, surprise, prediction error, entropy) included a constant and were estimated at the subject level. The significance of regression coefficients was estimated at the group level with t-tests. For multiple regressions, explanatory variables were z-scored so that regression coefficients can be compared between the variable of a given regression. Unless stated otherwise, all t-tests are two-tailed.

### Availability of data and code

The source data for all participants is available as **supplementary information**. The code to compute the ideal observers is available on GitHub: https://github.com/florentmeyniel/MinimalTransitionProbsModel

## ACKNOWLEDGEMENTS

We wish to thank Stefano Palminteri, Maxime Maheu, Peter Latham and Alexandre Pouget for useful comments on this work, and Bas Cornelissen for his help with Figure 3.

This work was supported by CEA (FM) and VSB Fund, Prins Bernhard Cultuurfonds and Institut français des Pays-Bas Descartes fellowships (MH).

## SUPPLEMENTARY INFORMATION

### SUPPLEMENTARY FIGURES AND LEGENDS

**Figure S1:**
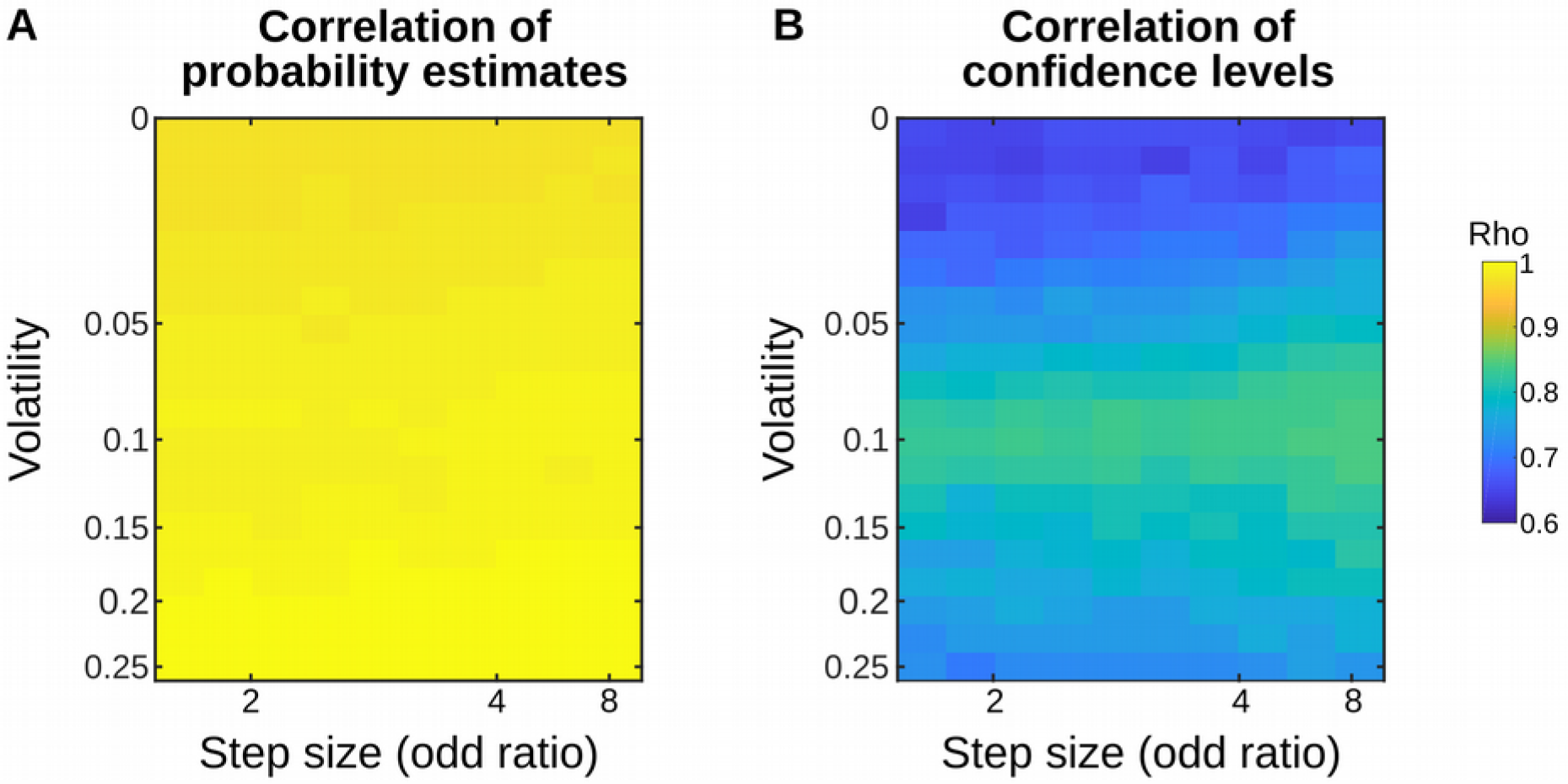
Correlation between the hierarchical and flat models is different for probability estimates and confidence levels. We ran simulations of our experiment (Fig. 2A) to assess which metric (probability estimates or confidence in those estimates) better distinguishes the hierarchical and flat learning models. For the sake of generality, we varied the volatility (probability of a change point in a sequence) and the step size of those changes (minimum fold change, in odd ratio, affecting the transition probabilities). For each combination of volatility and step size, we simulated 100 sequences to achieve stable results and we fit the single free parameter of each model (prior estimate of change point probability *p*_*c*_ in the hierarchical model; and leak factor *ω* in the flat model) onto the actual generative probabilities of the observed stimuli in the sequences. The resulting parameterized models therefore return their best possible estimate of the hidden regularities, in each volatility-step size condition. We then simulated new sequences (again, 100 per condition) to measure the correlation between (**A**) the estimated probabilities of stimuli between the two models, and (**B**) the correlation (Pearson’s rho) between the confidence (log-precision) that those models entertained in those estimates. The correlations indicate that probability estimates are nearly indistinguishable between the two models, whereas their confidence levels are more different. Note that the volatility level (0.013) and step size (4) used in the experiment ensure that confidence levels greatly differ between models. Those simulations used prior [1 1] for the flat model, but the results are qualitatively similar with prior [0 0] (see Methods).

**Figure S2:**
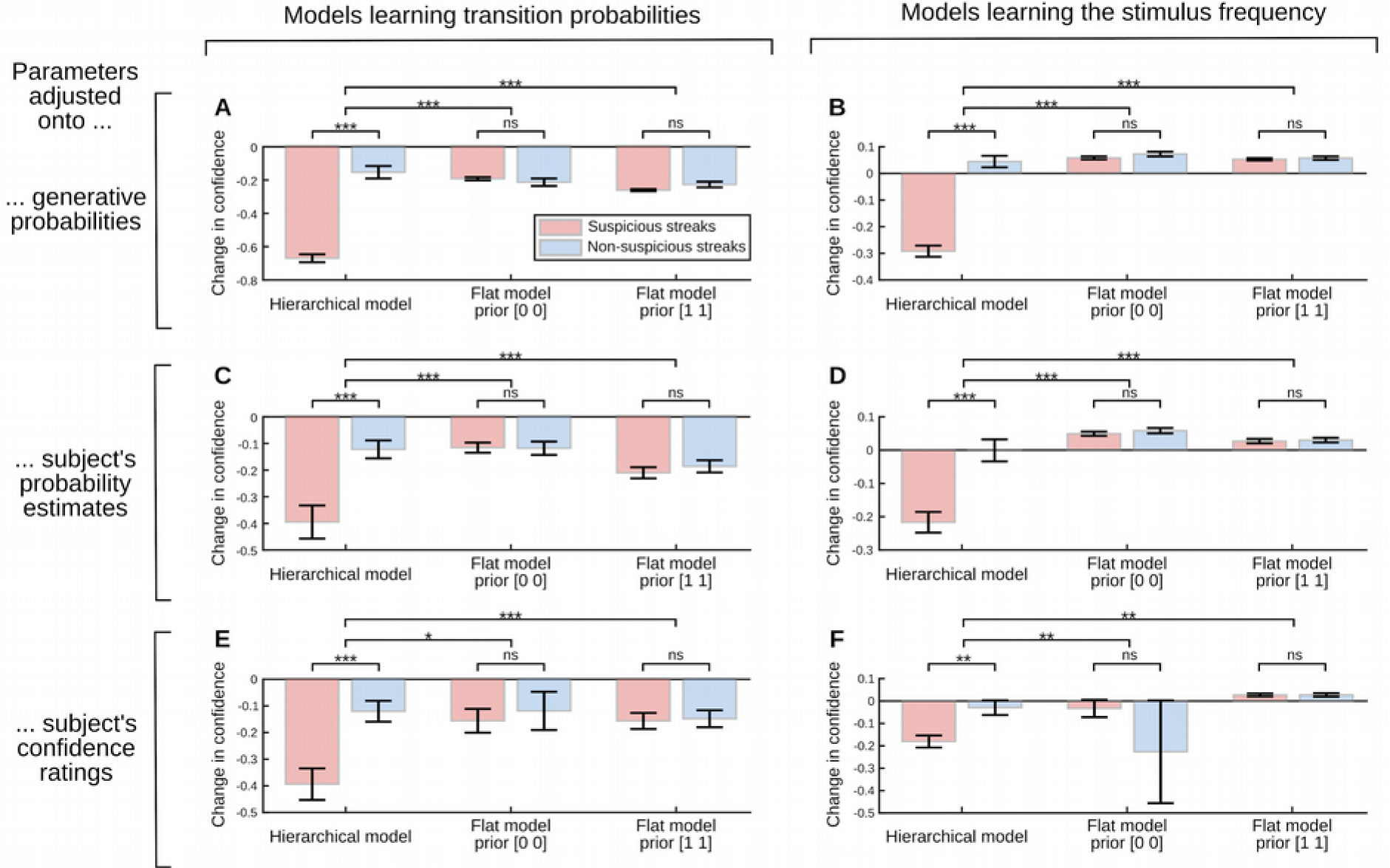
Theoretical predictions for various alternative models. Simulated changes in confidence around the target streaks in the main task. We consider three ideal observers models: the hierarchical model, the flat model with prior [0 0] and with prior [1 1]. Those models tracked either the transition probabilities between successive stimuli (**A, C, E**) or the frequency of stimuli (**B, D, F**). The free parameters of the models (the prior probability of change point p<_c_ in the hierarchical model; the leak factor ω in the flat model) were fitted following three procedures: so as to provide the best estimation of the actual generative probabilities of the sequences at all trials (**A, B**) or the answers of subjects at the moment of all questions regarding probability estimates (**C, D**) or their confidence ratings (**E, F**). Panels **A** and **B** therefore show the results of models that were optimized to solve the probability estimation task. By contrast, panels **C, D, E, F**, show the results of models that were optimized to be as close as possible to subjects, which can in principle deviate from **A** and **B**. Note that panel A corresponds to Fig. 3B, expanded with a new case (simulation of a flat model with priors [1 1]). None of the flat models in all plots shows an effect of streak type; some even predict increase (not decrease) in confidence (**B, D, F**). By contrast, a hierarchical model learning the stimulus frequency (**B, D, F**) seems more compatible with the subject’s data: they indeed predict an effect of streak type. One could therefore wonder whether subjects actually monitor the item frequency, instead of transition probabilities in the task. Several pieces of evidence argue against this possibility (see **Supplementary Results 3**). In addition, this possibility is incompatible with the results of the control experiment (Fig. 4B): a model that estimates a single statistic, namely the item frequency and the associated confidence, shows the same effect of streak type no matter whether change points are coupled between transition probabilities (main experiment) or uncoupled (control experiment). Data points are mean ± s.e.m of model predictions for each subject; significance levels correspond to p<0.05 (*), p<0.01 (**), p<0.001(***) in paired t-test, except for comparisons involving the flat model with prior [0 0] in **F** for which we used a Wilcoxon sign rank test because of an outlier point (see large error bar).

**Figure S3:**
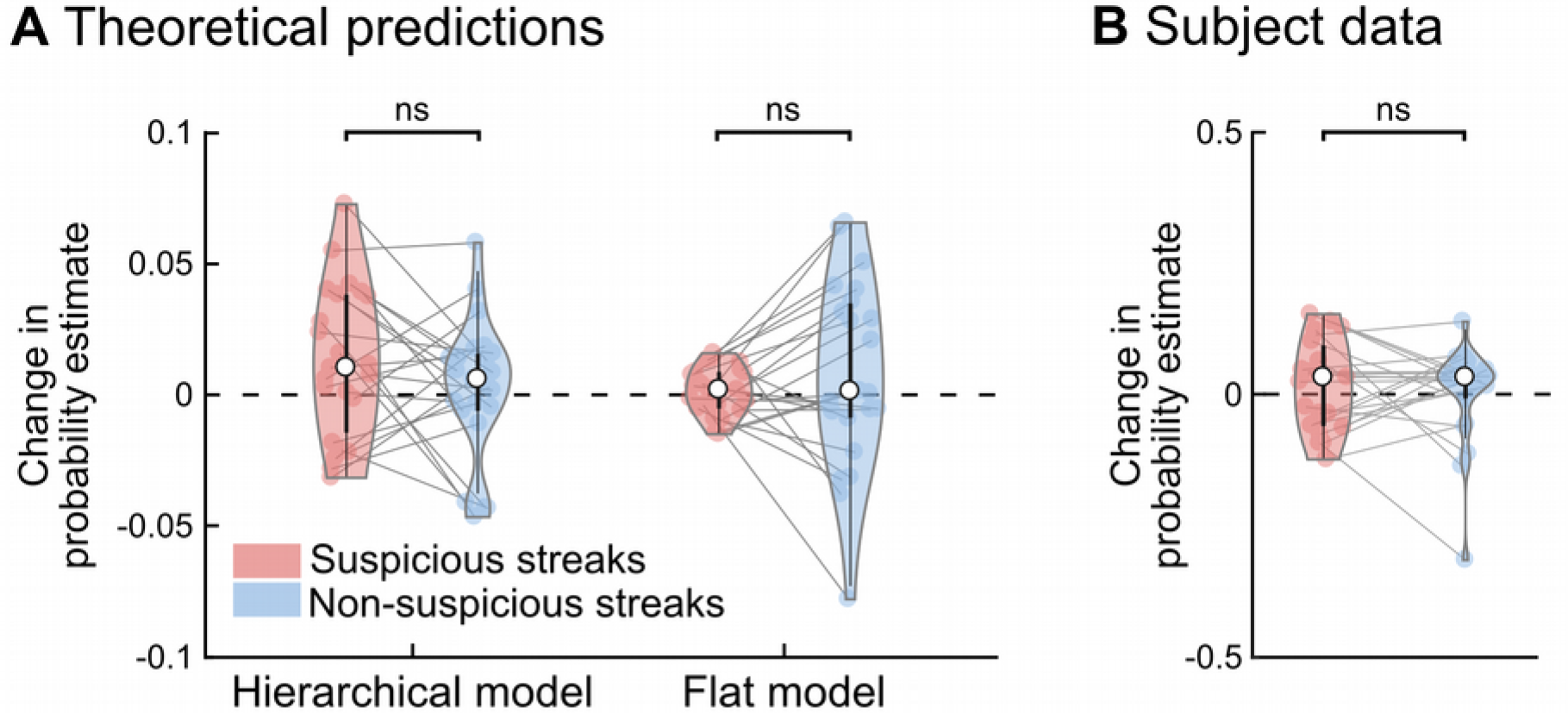
Probability estimates show no effect of streak type. This figure is analogous to Fig. 3B-C, excepted that it shows probability estimates reported at the moment of the pre/post streak questions, rather than the associated confidence levels. The error bars correspond to the inter-subject quartiles, distributions show subjects’ data; the significance level ‘ns’ corresponds to paired t-tests with p>0.15.

### SUPPLEMENTARY RESULTS

#### 1 Relation between the flat model and the delta rule

The equations of the flat model can be re-arranged so as to show the link with a leaky integrator, and hence, to a delta rule. For simplicity we derive those equations for the Bernoulli case (when one seeks to infer the frequency of items in a sequence), noting that the case of transition probabilities between successive items (p(A|A), p(B|B)) is nothing but the Bernoulli case when looking at each transition type separately (AA, BB).

Let’s recode the binary sequences as 1s and 0s, and estimate the probability P_1_(n) of observing a 1 after a sequence of observations y_1_, …, y_n_. P_1_(n) is the mean of a beta distribution, whose parameters are the (leaky) counts of observations and the prior count (cf. Eq. 3 and 5). Using the analytical solution for beta distributions, and recalling that the observation count is leaky, with exponential decay ω:

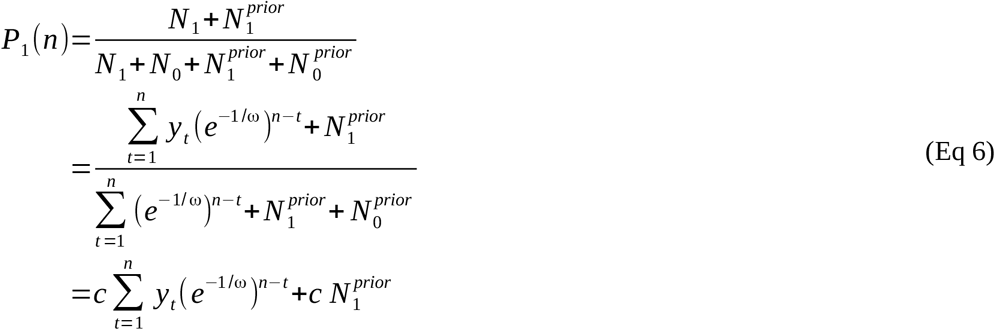

Where *c* can be approximated by a constant since e^−1/ω^<1 and *n* is typically large:

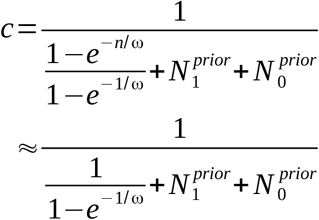

A delta-rule with learning rate α reads as follow:

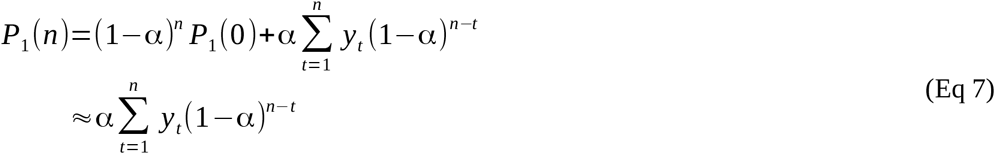

Note that for large *n*, the term before the sum vanishes since (1-α)<1, hence the approximation on the second line.

Comparison of Eq. 6 and 7 shows that the flat model and the delta rule are very similar, the only difference is that the leaky integration constantly adds a prior count in the flat model, whereas in the delta-rule, the impact of the starting point P_1_(0), which can be thought of as a prior, vanishes as more observations are accumulated. Stated differently, in the flat model, the inference progressively forgets about previous observations (like in the delta rule) and constantly factors in a prior about the estimated quantity (unlike the delta rule). Note that with N^prior^_1_ =N^prior^_0_ =0, both models become asymptotically identical for large *n*.

Computing the apparent learning rate (the ratio between update P_1_(n)-P_1_(n-1) and prediction error y_n_-P_1_(n-1) leading to this update) shows, with a bit of math, that for typical choices of ω, N^prior^_1_ >0 and N^prior^_0_ >0, the apparent learning rate increases whenever two consecutive observations are identical, and decreases whenever they differ. Considering that consecutive observations are more likely to differ after a change point in the underlying generative probability, the learning rate of the flat model typically increases, on average, immediately after change points (see Fig. 1).

#### 2 Robustness of the results

Each subject performed four blocks, two with auditory stimuli and two with visual stimuli. We tested the robustness of the linear relations between subject’s and optimal values (Fig. 2B-C) by testing them separately in each block type. We found that the results were replicated in each sensory modality. For probability estimates, in the auditory modality β=0.69±0.07 s.e.m., t_22_=9.27, p=4.7 10^−9^; in the visual modality β=0.64±0.06 s.e.m., t_22_=10.32, p=6.8 10^−10^. For confidence, in the auditory modality β=0.10±0.04 s.e.m., t_22_=2.85, p=0.009; in the visual modality β=0.09±0.03 sem, t_22_=2.81, p=0.010. Interestingly, the regression coefficients were correlated across subjects between modalities (probability estimates: ρ_23_=0.45, p=0.031; confidence ratings: ρ23=0.81, p=2.6 10^−6^), suggesting that inference in this task operates at an abstract, amodal level.

We further tested the robustness of the correlation between subject’s and optimal values by restricting the regression analysis to a subset of data points, namely, the questions that surround the target streaks (Fig 3A). The significant correlations were replicated on this subset of data for both probability estimates (β=0.57±0.07 s.e.m., t_22_=7.6, p=1.4 10^−7^) and confidence ratings (β=0.12±0.05 s.e.m., t_22_=2.6, p=0.017).

We also tested the robustness of our central analysis of the effect of streak type on the change in confidence. In the main text, we report a dichotomy between suspicious and non-suspicious streaks, but in reality the ‘suspiciousness’ of a streak is a matter of degree: the more a streak arouses the suspicion of a change, the larger the decrease in confidence. We therefore regressed the subject’s changes in confidence onto the hierarchical model’s changes in confidence across all streaks. In order to test whether the hierarchical model or the flat model provides a better account of the subjects’ data, we also include the change in confidence of the flat model as a competing explanatory variable in the multiple regression. Regression coefficients were significant for the hierarchical model (β=0.07±0.02 s.e.m., t_22_=3.7, p=0.001), but not for the flat model (β=0.01± 0.01 s.e.m., t_22_=0.8, p=0.44), and the regression coefficients of the hierarchical model were significantly larger than those of the flat model (paired difference of βs=0.06±0.03 s.e.m., t_22_=2.3, p=0.031), indicating that the hierarchical model provides a significantly better account of the subjects’ data.

#### 3 Normative properties of confidence reports

In the ideal observer model, the answers to questions #1 and #2 (probability estimate and confidence rating) are different readouts of the same posterior distribution, namely its mean and log-precision. If the subjects’ answers to those questions also derive from the same inference process, then we expect that subjects who are closer to the optimal probability estimates are also those who are closer to the optimal confidence ratings. We therefore tested whether the linear regression coefficients (βs) linking subjects and the optimal hierarchical model were correlated between probability estimates and confidence ratings. The between-subject correlations was indeed significant: ρ_23_=0.53, p=0.009.

We tested for further normative properties of the subject’s confidence ratings. Several factors, notably factors pertaining to first-order estimates, are expected to impact confidence ratings in this task from a normative viewpoint. We first show that those factors indeed impact the optimal confidence levels at the moments of questions during the task, and then report a similar analysis for the subject’s confidence ratings. Optimal confidence levels were entered into a multiple regression model which included the estimated probability itself, the entropy of this probability (which quantifies the estimated unpredictability of the next stimulus, it culminates when the estimated probability is 0.5), and the extent to which the current observation deviates from the previous estimate, as quantified by the surprise (negative log likelihood of the observations, (Shannon, 1948)) and the prediction error (one minus the likelihood of the current observation). Optimal confidence was lower when the estimated entropy was higher (β=−0.051±0.008, t_22_=−6.1, p=4.2 10^−6^), lower when surprise was larger (β=−0.142±0.026, t_22_=−5.4, p=1.8 10^−5^) and when prediction error was larger (β=−0.128±0.030, t_22_=−4.2, p=3.5 10^−4^).

To analyze subjects’ confidence ratings, we added other explanatory variables to this multiple linear regression model, which correspond to subject’s estimates: the subject’s probability estimate, and the entropy corresponding to this estimate. Note that questions are asked only occasionally, so that we don’t know the probability estimate of the subject at the *previous trial*, and therefore, we cannot compute the subject’s surprise and prediction error elicited by the last observation. Subject’s confidence was lower when the entropy of his estimate was higher (β=−0.125±0.009, t_22_=−14.5, p=1.0 10 ^−12^) and when the optimal surprise level was higher (β=−0.090±0.022, t_22_=−4.1, p=5.0 10^−4^).

Another aspect of subjects’ accuracy is that their report of confidence is specific to the relevant statistics. In the experiment, subjects monitor two transition probabilities, there are therefore two confidence levels, each being attached to one transition type. Questions asked subjects to estimate the likelihood of the next stimulus, which depends on only one of the two transition probabilities: the one that is relevant given the identity of the previous stimulus at the moment of the question. We estimated a multiple linear regression in which the subject’s confidence ratings was regressed onto both the optimal relevant confidence levels, and the optimal irrelevant confidence level (those attached to the irrelevant transition). The regression coefficients corresponding to the relevant confidence levels were significant (β=0.039±0.014 s.e.m., t_22_= 2.8, p=0.011), those for the irrelevant confidence were not (β=0.001±0.008 s.e.m., t_22_=0.1, p=0.89) and the difference between the two was significant (paired difference of βs=0.038±0.018 s.e.m., t_22_=2.0, p=0.027, one-tailed test). A different multiple linear regression indicates, in addition, that confidence ratings are selectively modulated by the (optimal) entropy of the relevant transition probability, as opposed to the irrelevant one (paired difference of βs=−0.020±0.007 s.e.m., t_22_;−2.9, p=0.008). Optimal confidence levels show the same effect: replacing the subjects’ confidence ratings in this latter regression with the optimal confidence levels also reveals a significant difference (paired difference of βs=−0.050±0.016, t_22_=−3.2, p=0.0045). By contrast, such a difference is not observed when replacing the subjects’ confidence ratings with the optimal confidence of a model that monitors solely the frequency of items (paired difference of βs=0.005±0.008, t_22_=0.6, p=0.54). Together, those results indicate that subjects reported specifically the confidence attached to the transition probability relevant at the moment of the question.

#### 4 Theoretical effects on the apparent learning rate

We cannot assess the apparent learning rate of subjects on a trial-by-trial basis here since it would require that subjects report their first-order estimates on every trial, whereas they did it only occasionally. However, we can run such an analysis on our simulated models. We found a specific effect of streak type on the apparent learning rate of the hierarchical model, which increased more after suspicious streaks than non-suspicious ones (0.11±0.01 s.e.m., p=2.9 10^−14^, t_22_=17.2), there was no difference in the flat model (−0.0023±0.0022 s.e.m., p=0.3, t_22_=−1.1) and the difference between models was significant (paired difference of differences, −0.11±0.01 s.e.m., p=1.4 10^−14^, t_22_=−17.9). This effect of streak type in the hierarchical model for uncoupled change points was no longer observed in the control task with uncoupled change points (−0.001±0.005 s.e.m., p=0.85, t_20_=−0.2). In other words, the apparent learning rate passes the test we propose to detect the use of a hierarchical model.

### SUPPLEMENTARY DATA

We provide the raw data as a Matlab data file: *HierarchyTasks_FullRawDataSet.mat*. This data file can be read with Matlab, or a freely available software such as GNU Octave or Python. The file contains three cell variables: one for the subjects included in the main task, one for the subjects excluded from the main task, and one for the subjects (all included) in the control task. Each element of a cell corresponds to one subject, and the data are presented as a matrix. Each row is a trial, and the columns should be read as follows:

1. Sensory modality (“1” for visual, “0” for auditory)
2. Block number (1 to 4)
3. Observed binary sequence, coded as “1” and “2”
4. Generative probability of observing “1” when the previous stimulus is “2”
5. Generative probability of observing “2” when the previous stimulus is “1”
6. Subject’s estimate of the probability of receiving “1” on the next trial, from 0 to 1 (with NaN when no question is asked)
7. Subject’s confidence about the estimated probability, from 0 to 1 (with NaN when no question is asked)
8. Reaction times (s) for the probability report (with NaN when no question is asked)
9. Reaction time (s) for the confidence report (with NaN when no question is asked)

